# Lipid droplet isolation as a novel platform for spectroscopic investigation of cargo modifications

**DOI:** 10.64898/2026.01.30.702822

**Authors:** Karolina Chrabąszcz, Agnieszka Panek, Katarzyna Pogoda

## Abstract

Lipid droplets (LDs) are dynamic organelles involved in metabolic regulation and cellular stress responses, yet their biochemical heterogeneity and treatment-dependent remodeling in the context of radiotherapy remain poorly understood. Here, we present the first label-free Raman spectroscopic analysis of isolated lipid droplets (iLDs) from normal Schwann cells and malignant peripheral nerve sheath tumor (MPNST) cells subjected to cannabidiol (CBD) treatment, ionizing radiation, and their combination. Raman spectroscopy revealed pronounced chemical heterogeneity of iLDs both between and within cell types, reflecting differences in acyl chain organization, conformational order, and lipid class composition. CBD treatment induced substantial lipid remodeling in Schwann cells, giving rise to multiple iLD subpopulations, whereas MPNST cells exhibited a more constrained response. Irradiation altered lipid droplet heterogeneity in a cell-type-dependent manner, while combined CBD treatment and irradiation induced characteristic alterations in LD cargo that differed markedly between Schwann and MPNST cells, highlighting lipid droplets as sensitive reporters of metabolic reprogramming and stress adaptation. Overall, these findings establish Raman-based lipid droplet profiling as a powerful approach for resolving treatment-specific metabolic remodeling at the suborganelle level and provide new insight into lipid-mediated mechanisms underlying radiosensitization in cancer cells.

## 1. Introduction

Cellular adaptation to physiological conditions and external stress requires rapid and coordinated reorganization of metabolic resources.^[1],[2]^ Among the major classes of biomolecules, lipids occupy a central position due to their dual roles as structural components and active regulators of cellular signaling.^[3]^ Beyond forming biological membranes and serving as energy reservoirs, lipids participate directly in redox balance, stress sensing, and modulation of intracellular signaling pathways.^[4]^ Consequently, perturbations in lipid composition, organization, and turnover have profound effects on cellular homeostasis and are increasingly implicated in pathological conditions, including cancer, metabolic disorders, and neurodegenerative diseases.^[5]^

In tumor cells, lipid metabolism is frequently reprogrammed to support survival under adverse conditions such as nutrient limitation, oxidative stress, and therapeutic pressure.^[6],[7]^ A prominent manifestation of this metabolic rewiring is the altered formation, accumulation, and composition of lipid droplets (LDs), which function as dynamic lipid storage organelles and stress-adaptive hubs. Increased LD abundance has been reported across multiple cancer types and has been linked to enhanced resistance to ionizing radiation.^[8]^ By buffering oxidative stress, regulating lipid availability for membrane repair, and interacting with iron-dependent metabolic pathways, LDs contribute to cellular mechanisms that promote survival following radiation-induced damage. Despite growing recognition of these roles, the chemical nature of LD cargo remodeling associated with radioresistance remains poorly defined.

Current insights into LD biology are derived predominantly from fluorescence microscopy and bulk lipidomic analyses.^[9]^,^[10]^,^[11]^ While these approaches provide valuable information on LD abundance, spatial distribution, and overall lipid composition, they offer limited access to intrinsic chemical features such as lipid unsaturation, acyl chain ordering, or therapy-induced structural rearrangements within LD cargo. Moreover, extensive sample processing, labeling, and extraction steps can obscure subtle yet biologically meaningful molecular changes. These limitations underscore the need for analytical strategies capable of probing lipid chemistry directly and with minimal perturbation.

Vibrational spectroscopic techniques, and Raman spectroscopy in particular, provide a label-free and non-destructive means of interrogating molecular composition based on intrinsic vibrational signatures.^[12]^,^[13]^ Raman spectroscopy is especially well suited for lipid analysis, enabling sensitive detection of acyl chain saturation, conformational order, chain length, cis– trans isomerism of unsaturated bonds, membrane fluidity and phase state, lipid oxidation, and compositional differences among major lipid classes, as well as spatially resolved mapping of lipid heterogeneity at the cellular level. However, its application to lipid droplets has largely been restricted to in situ cellular measurements.^[14]^,^[13]^,^[5]^ Systematic Raman spectroscopic analysis of isolated LDs has remained challenging due to the lack of isolation strategies that preserve native chemical properties while ensuring compatibility with spectroscopic requirements.

Here, we introduce a lipid droplet isolation platform specifically tailored for Raman spectroscopic investigation and apply it to a biologically and clinically relevant model system. Lipid droplets were isolated from normal human Schwann cells and malignant peripheral nerve sheath tumor (MPNST) cells subjected to defined therapeutic conditions, including untreated controls, cannabidiol (CBD) treatment, ionizing radiation, and sequential CBD pre-treatment followed by irradiation (Figure 1). This experimental design enables direct comparison of LD cargo chemistry between non-malignant and malignant cells, as well as systematic assessment of therapy-induced lipid remodeling.

**Figure 1.**
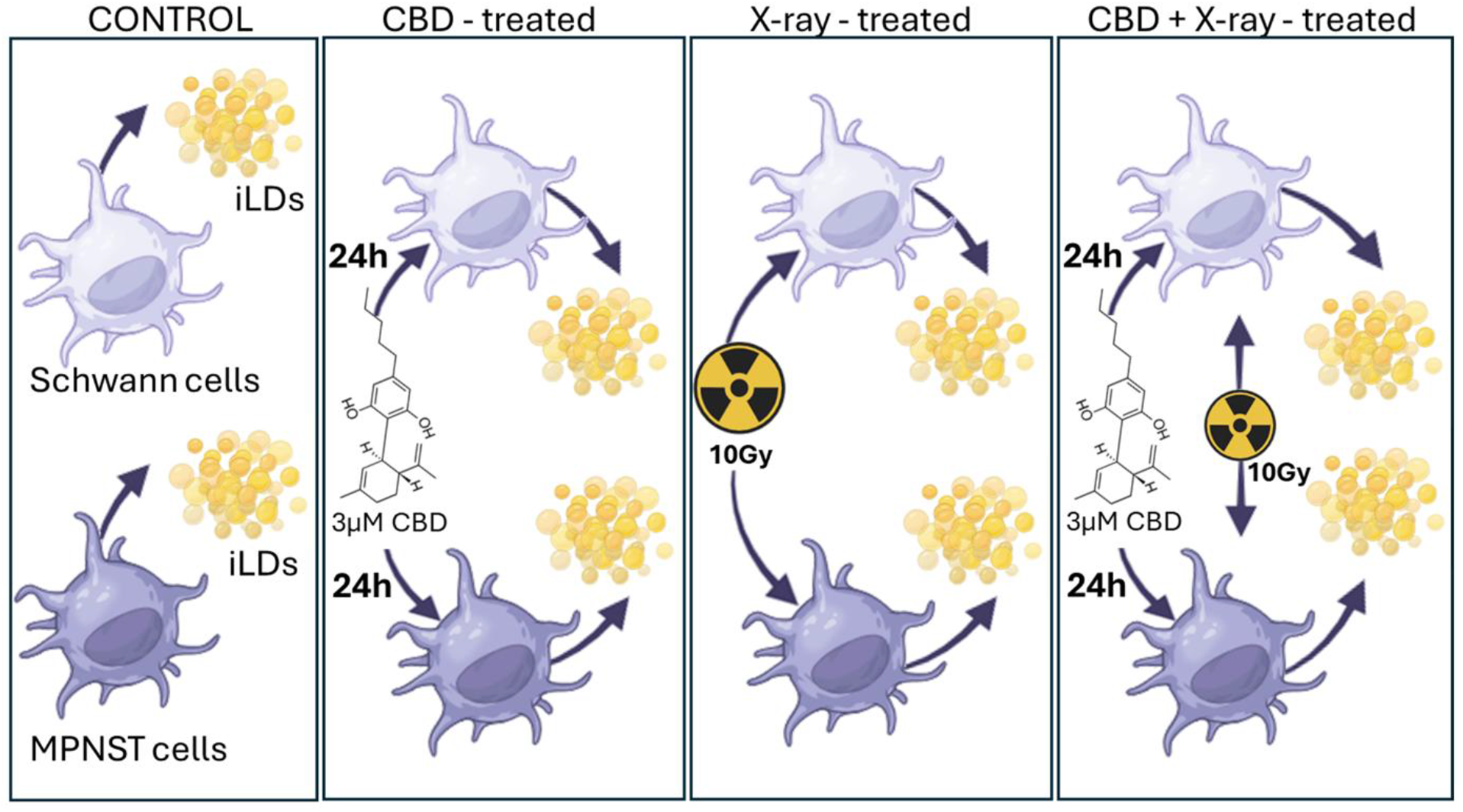
Schematic overview of experimental treatment conditions applied to Schwann and MPNST cells followed by the lipid droplet isolation (iLD). (From left) Lipid droplets were isolated from untreated cells, following cannabidiol (CBD, 3 µM, 24 h) treatment, X-ray irradiation (10 Gy), or combined CBD pre-treatment followed by irradiation.

Using Raman spectroscopy, we demonstrate that isolated lipid droplets retain distinct and condition-specific vibrational fingerprints, reflecting changes in lipid unsaturation, molecular packing, and overall chemical organization. Together, our results establish LD isolation coupled with Raman spectroscopy as a novel platform for label-free investigation of lipid droplet cargo modifications and provide new insights into lipid metabolic responses associated with cancer and therapeutic stress.

## 2. Results and discussion

### 2.1. Spectroscopic validation of lipid droplet isolation

To ensure that the Raman spectroscopic signals analyzed in this study originated specifically from isolated lipid droplets, a spectroscopic validation strategy based on deuterated palmitic acid (d_31_-palmitic acid, d_31_-PA) labeling was employed (Figure 2). Excess fatty acids are preferentially sequestered into lipid droplets, and incorporation of d_31_-PA was therefore expected to result in selective accumulation within these organelles.^[15]^ Importantly, C–D vibrational modes occur in the Raman silent region, providing an unambiguous spectral marker for lipid droplet–associated lipids and enabling their discrimination from endogenous cellular components.^[16]^ Because deuterated compounds may exhibit cytotoxic effects at elevated concentrations, MTS viability assays were first performed to determine a suitable labeling concentration (Figure 2A). These results revealed a concentration-dependent decrease in cell viability, with concentrations above 10 µM exerting a more pronounced effect on malignant MPNST cells than on normal Schwann cells. Based on these findings, and consistent with our previous demonstrations of Raman sensitivity to micromolar intracellular probe concentrations, 10 µM d_31_-PA was selected for all subsequent experiments.^[13]^ The Raman spectrum of the pure d_31_-PA standard confirms the characteristic isotopic shift of C–H stretching vibrations (2800-3000 cm^−1^) to the C–D region (2000-2300 cm^−1^), providing a distinct and interference-free spectral signature (Figure 2B).^[16]^

**Figure 2.**
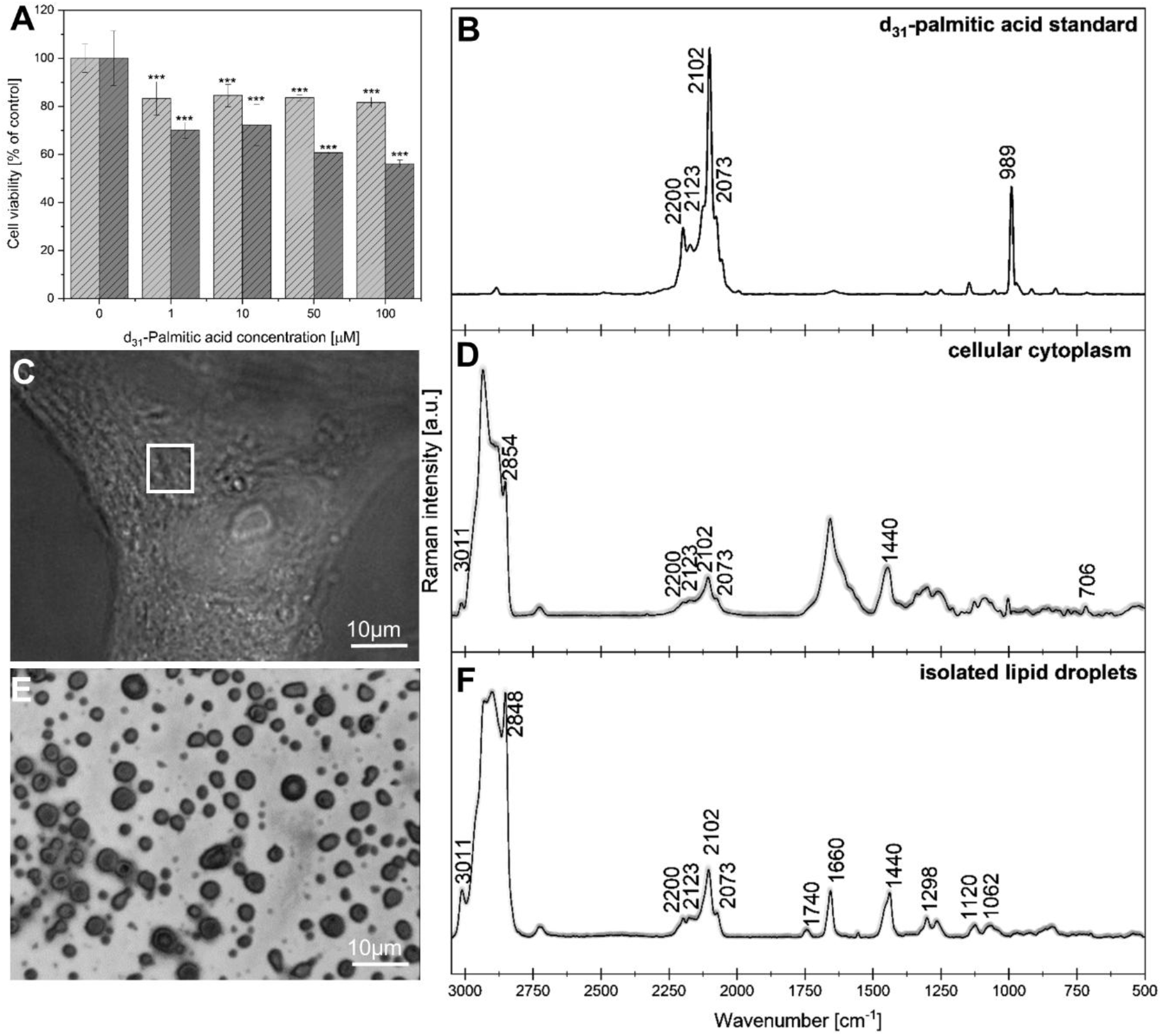
Results of the MTS assay performed on Schwann (light grey) and MPNST (dark grey) cell lines treated with increasing concentrations of acid d_31_-PA (A). Raman spectrum of the pure d_31_-PA standard (B). Brightfield image of a cell treated with 10 µM d_31_-PA with the region selected for Raman imaging indicated (C) together with the corresponding Raman spectrum acquired from the cytoplasm showing accumulation of d_31_-PA (D). Brightfield image of dried iLDs (E) with representative Raman spectra confirming their biochemical composition and the presence of incorporated d_31_-PA (F).

Following cellular incubation, Raman imaging of the cytoplasmic region revealed the presence of d_31_-PA-associated signals even in cases where lipid droplets were not readily discernible in brightfield microscopy (Figure 2C). Raman spectra extracted from these regions displayed a composite biochemical signature, including lipid-associated bands such as the CH_2_ stretching mode at ∼2850 cm^−1^, while also containing contributions from other cellular constituents (Figure 2D). These observations confirm intracellular accumulation of d_31_-PA but underscore the spectral complexity inherent to whole-cell measurements. Subsequent isolation of lipid droplets from d_31_-PA-treated cells, performed as described in the *Materials and Methods* section, enabled direct spectroscopic interrogation of lipid droplet cargo (Figure 2E,F). Brightfield imaging of isolated lipid droplets (iLDs) revealed the presence of larger agglomerates rather than individual droplets, likely resulting from necessary rinsing and drying steps required to remove buffer components with strong Raman signatures (*Materials and Methods*, Figure 5). Despite this aggregation, Raman spectra acquired from iLDs exhibited marked differences compared to spectra collected from intact cells (Figure 2E, F). In particular, the appearance of prominent bands at ∼1740 cm^−1^ and ∼1660 cm^−1^, attributed to ester carbonyl (C=O) and C=C stretching vibrations of cholesteryl esters, respectively, reflects the preferential storage of excess lipids in esterified forms within lipid droplets.^[16]^ Moreover, a strong and well-resolved C–D signal in the 2200-2073 cm^−1^ region confirms the presence of incorporated d_31_-PA within the isolated droplets. Notably, no spectral contributions from isolation buffers (Figure 5) were detected, indicating effective removal of extraneous components and preservation of lipid droplet chemical integrity.

These results confirm that the applied isolation protocol yields lipid droplet-enriched fractions with preserved and spectroscopically distinguishable lipid signatures. The use of d_31_-PA provides a robust and selective validation strategy, establishing a solid foundation for subsequent Raman-based analysis of lipid droplet heterogeneity and treatment-induced remodeling.

### 2.2. Lipid droplet heterogeneity revealed by Raman spectral signatures

Representative Raman spectra of iLDs derived from normal Schwann cells and MPNST cells revealed pronounced spectral heterogeneity both between and within the two cell types (Figure 3). Although all spectra were dominated by lipid-associated vibrational features, marked differences in spectral complexity and band intensities indicated the presence of chemically distinct lipid droplet subpopulations. In the high-wavenumber region, intense CH_2_ stretching bands at ∼2846-2850 cm^−1^ and ∼2880-2887 cm^−1^ were observed across all iLD spectra, reflecting the abundance of lipid acyl chains. Variations in relative band intensities among spectra originating from the same cell type point to the coexistence of multiple lipid droplet populations. In particular, differences in the CH_2_ scissoring region and in C–C stretching modes reflect variability in acyl chain ordering, lipid saturation, and backbone conformation, underscoring the intrinsic compositional diversity of lipid droplet cargo.Importantly, this heterogeneity was not limited to differences between Schwann and MPNST cells. Within Schwann cells, one subclass of iLDs exhibited a richer spectral profile, characterized by additional bands at ∼1337 cm^−1^, ∼1167-1152 cm^−1^, and in the low-wavenumber region (∼840-794 cm^−1^), indicative of contributions from multiple lipid classes and structurally diverse lipid species (Figure 3A). In contrast, another Schwann cell iLD subpopulation displayed a simplified Raman signature dominated by CH_2_ scissoring (∼1440-1462 cm^−1^) and C–C stretching modes (∼1128-1060 cm^−1^), consistent with a more uniform, fatty acid-like lipid composition. Similarly, both identified subclasses of MPNST-derived iLDs exhibited comparatively simplified Raman profiles dominated by CH_2_ scissoring and C–C stretching modes, suggesting preferential enrichment in specific lipid subclasses or fatty acid–rich compositions (Figure 3C, D).

**Figure 3.**
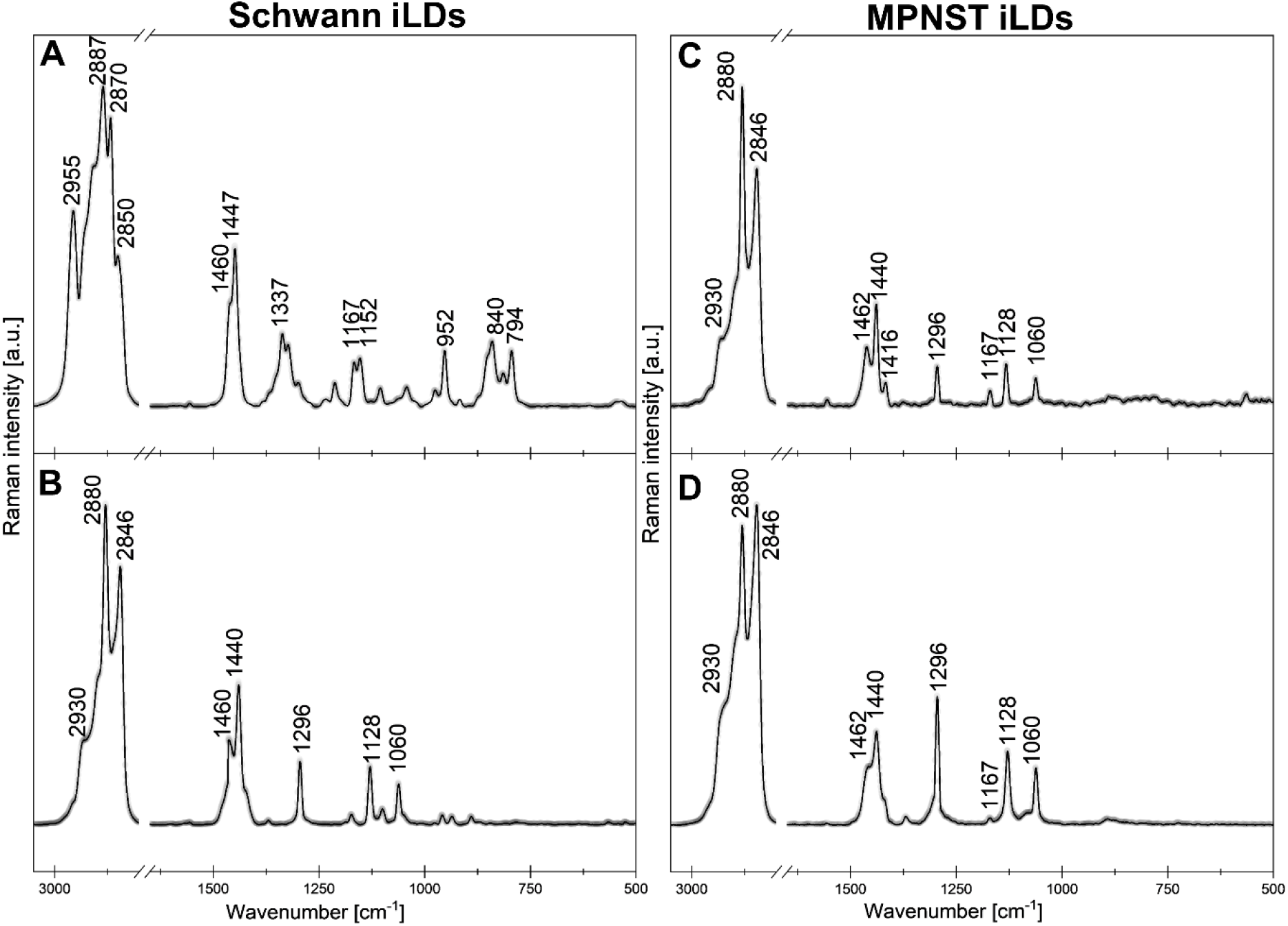
Representative Raman spectra of isolated lipid droplets (iLDs) obtained from Schwann cells (A, B) and MPNST cells (C, D). Variations in band presence and relative intensities reflect chemical heterogeneity of lipid droplet cargo both between cell types and within individual iLD populations, indicating the coexistence of spectrally distinct lipid droplet subtypes.

Together, these results demonstrate that isolated lipid droplets represent chemically heterogeneous organelles rather than uniform lipid reservoirs, which have more complex spectral pattern for Schwann cells subpopulation. The fact that distinct lipid droplet groups are already present under basal conditions strongly suggests that they may exhibit differential metabolic responsiveness and undergo selective remodeling upon metabolic stress or therapeutic perturbations.

### 2.3. Therapy-induced diversification of lipid droplet spectral phenotypes

Based on our previous studies demonstrating that cannabidiol (CBD) induces lipid remodeling that enhances the radiosensitivity of radioresistant MPNST cell lines^[17]^, we sought to elucidate the lipid-related mechanisms underlying this effect by investigating how different treatment strategies influence the biochemical composition of intracellular lipid droplets (iLDs) (Figure 4).

**Figure 4.**
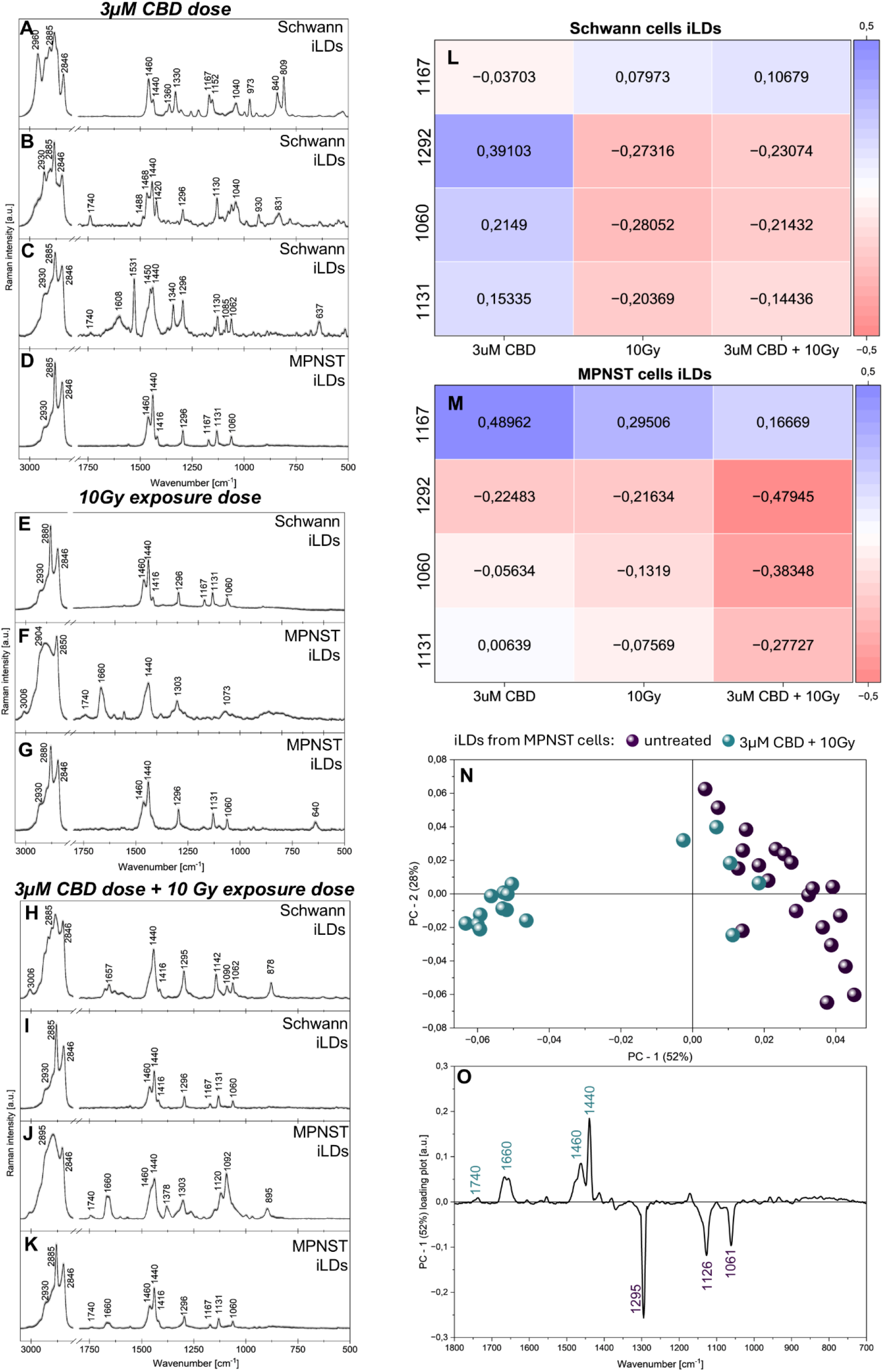
Raman spectra of iLDs from Schwann and MPNST cells following treatment with 3 µM cannabidiol (CBD), 10 Gy irradiation, or combined CBD and irradiation (A–K). Heatmaps show treatment-dependent changes in selected Raman bands associated with lipid chain conformation and packing in Schwann cell–derived (L) and MPNST-derived (M) iLDs. Principal component analysis (PCA) of selected MPNST iLD spectra (N) reveals clear separation between untreated and combined treatment conditions along PC1 (52% variance), with corresponding loadings (O) indicating coordinated remodeling of lipid droplet composition and acyl chain organization.

**Figure 5.**
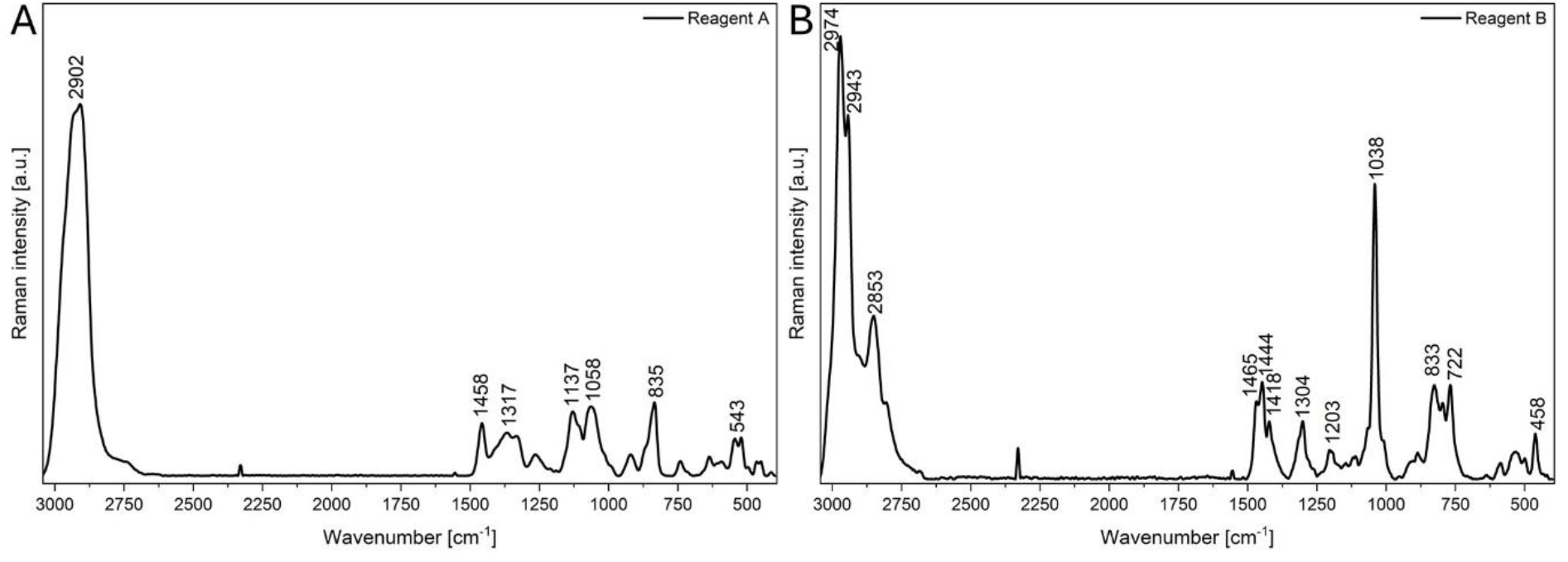
Raman spectra of reagent A (A) and reagent B (B) used during lipid droplets isolation.

First, the effect of CBD alone was examined in both cell lines using a concentration of 3 µM, previously identified as biologically effective.^[17]^ Raman spectral analysis revealed marked differences in lipid droplet heterogeneity between normal Schwann cells and MPNST cells (Figure 4A-D). In Schwann cells, three distinct iLD subpopulations were identified, indicating pronounced lipid remodeling in response to CBD. In contrast, only a single lipid droplet population was detected in MPNST cells, suggesting a more limited lipid response to CBD treatment. The first Schwann cell iLD subpopulation was dominated by CH_2_ stretching vibrations (∼2850-2885 cm^−1^), CH_2_ scissoring modes (∼1440-1460 cm^−1^), and C–C stretching bands at ∼1130 and ∼1167 cm^−1^ associated with all-trans conformations, consistent with neutral lipids characterized by highly ordered acyl chains (Figure 4A).A second subpopulation exhibited an additional pronounced ester carbonyl band at ∼1740 cm^−1^ together with the CH_2_ twisting mode at ∼1296 cm^−1^, indicative of a mixed lipid composition comprising neutral lipids (Figure 4B). The third iLD subpopulation displayed ester carbonyl (∼1740 cm^−1^), phosphate-related vibrations around ∼1085 cm^−1^, enhanced C–C stretching at ∼1060 cm^−1^ associated with gauche conformations, and ∼1531 cm^−1^ assigned to amide II modes, respectively, suggesting a membrane-associated lipid environment enriched in phospholipids and/or sphingolipids, potentially coexisting with membrane-associated proteins (Figure 4C). In contrast, CBD-treated MPNST cells exhibited a single iLD population resembling the control lipid profile, albeit with reduced intensities of bands at ∼1296, ∼1167, ∼1131, and ∼1060 cm^−1^. These bands originate from CH_2_ twisting and C–C stretching vibrations of lipid acyl chains, with ∼1167 and ∼1131 cm^−1^ associated with all-trans conformations, ∼1060 cm^−1^ reflecting gauche conformations linked to increased chain disorder and membrane fluidity, and ∼1296 cm^−1^ sensitive to acyl chain packing and conformational order. ^[18],[16],[19]^

Upon irradiation with a 10 Gy dose, reduced lipid droplet heterogeneity was observed in both cell lines (Figure 4E-G). In Schwann cells, iLDs displayed fatty acid-like spectral profiles with altered intensity ratios of the 1460, 1440, and 1416 cm^−1^ bands, corresponding to CH_2_ scissoring and asymmetric and symmetric CH_3_ deformation modes, indicating changes in acyl chain organization compared to untreated cells. In irradiated MPNST cells, two lipid droplet populations emerged, one resembling a cholesteryl ester-like profile and the other exhibiting a fatty acid-like spectral signature, reflecting altered lipid composition and acyl chain organization.

Following combined CBD incubation and irradiation (Figure 1), two distinct iLD spectral populations were identified in both Schwann cells (Figure 4H,I) and MPNST cells (Figure 4J,K). In Schwann cells, the two populations differed mainly in CH_2_-related band intensities and acyl chain order, consistent with moderate heterogeneity within fatty acid–like neutral lipid droplets. In contrast, iLDs isolated from MPNST cells showed more pronounced spectral divergence, sharing common ester carbonyl (∼1740 cm^−1^) and unsaturated C=C (∼1660 cm^−1^) bands while differing substantially in CH_2_ stretching and deformation modes and in trans/gauche-related C–C vibrations. This pattern indicates a stronger treatment-induced remodeling of lipid composition and organization in cancer cells, involving the coexistence of ordered neutral lipids and a second population enriched in ester-containing and membrane-associated lipid species.

Given the high spectral complexity and heterogeneity of iLDs, subsequent quantitative analysis focused on Raman bands consistently observed in at least one lipid droplet population within each experimental condition. Bands at ∼1167, ∼1292, ∼1060, and ∼1031 cm^−1^ were selected, as they report on lipid chain packing, conformational rearrangements, relative ordering, and compositional differences between fatty acid–like and more complex lipid species. Changes relative to untreated controls were visualized as heatmaps (Figure 4L,M).

Such analysis revealed distinct treatment-dependent lipid responses in Schwann and MPNST cells. In Schwann cell-derived iLDs (Figure 4L), CBD treatment was associated with positive correlations for the ∼1292, ∼1060, and ∼1131 cm^−1^ bands, indicative of increased CH_2_ twisting and enhanced contributions from gauche-related C–C stretching modes, consistent with moderate conformational disorder and reduced lipid chain packing. In contrast, irradiation alone and combined CBD plus irradiation were characterized by negative correlations for these bands and relatively increased contributions of the ∼1167 cm^−1^ band, suggesting a shift toward more ordered, trans-rich acyl chain conformations. In MPNST-derived iLDs (Figure 4M), CBD treatment alone induced a strong positive correlation of the ∼1167 cm^−1^ band together with negative correlations of the ∼1292 and ∼1060 cm^−1^ bands, consistent with increased lipid chain ordering and reduced conformational flexibility. Notably, combined CBD and irradiation resulted in pronounced negative correlations for the ∼1292, ∼1060, and ∼1131 cm^−1^ bands, indicating substantial suppression of gauche conformations and marked remodeling of lipid chain organization. Overall, these patterns demonstrate a differential regulation of lipid droplet composition and acyl chain ordering in response to CBD and irradiation, with cancer cells exhibiting a stronger and more coordinated shift toward ordered lipid structures under combined treatment conditions. This effect is consistent with the lipophilic nature of CBD and its ability to intercalate into lipid membranes and droplets, thereby perturbing lipid–lipid and lipid– protein interactions and altering the functional consequences of increased lipid ordering under irradiation-induced stress.

As the most pronounced alterations were observed for MPNST-derived iLDs under treatment combination, heatmap-based analysis was further supported by principal component analysis (PCA) for this condition (Figure 4N,O). PCA revealed clear separation between untreated iLDs and iLDs derived frm cells under combined CBD with irradiation strategy, primarily along PC1, accounting for 52% of the total variance. The corresponding PC1 loading plot showed strong positive contributions from bands at ∼1440-1460, ∼1660, and ∼1740 cm^−1^ and negative loadings at ∼1295, ∼1126, and ∼1061 cm^−1^, in full agreement with the heatmap results.

Together, these analyses demonstrate that combined CBD and irradiation induce a coherent and distinct remodeling of lipid droplet composition and acyl chain organization in MPNST cells, consistent with the formation of more ordered yet functionally altered lipid structures.

## Conclusions

In this study, we provide the first comprehensive Raman spectroscopic characterization of treatment-induced biochemical remodeling of intracellular lipid droplets in normal Schwann cells and radioresistant MPNST cells. By combining lipid droplet isolation with label-free Raman spectroscopy and multivariate analysis, we demonstrate that iLDs are highly dynamic and chemically heterogeneous organelles whose composition and acyl chain organization are selectively reshaped by cannabidiol treatment, irradiation, and their combination.

Importantly, our results reveal that distinct lipid droplet subpopulations are already present under basal conditions and undergo divergent remodeling pathways depending on the applied therapeutic strategy. The observed treatment-specific changes in lipid chain packing, conformational order, and lipid class composition highlight lipid droplets as sensitive reporters of cellular metabolic state and stress responses. Notably, cancer cells exhibited a more coordinated and pronounced lipid remodeling under combined CBD and irradiation, underscoring a previously unexplored link between lipid droplet organization and radiosensitization.

These findings establish Raman-based lipid droplet profiling as a powerful and forward-looking approach for studying lipid metabolism in the context of cancer therapy. The ability to resolve treatment-dependent changes in iLD composition at the subpopulation level opens new avenues for understanding adaptive lipid remodeling mechanisms and for identifying lipid-based biomarkers of therapeutic response. As such, this strategy holds significant potential for future applications in radiobiology, metabolic targeting, and personalized cancer treatment.

## Materials and Methods

### Cell Culture

Normal human Schwann cells (hTERT NF1 ipnNF95.11c) and malignant peripheral nerve sheath tumor cells (MPNST; sNF02.2) were cultured in Dulbecco’s Modified Eagle Medium (DMEM) supplemented with 10% fetal bovine serum (FBS) and 100 U mL^−1^ penicillin– streptomycin. For MPNST cells, the medium was additionally supplemented with L-glutamine. Cells were maintained in 75 cm^2^ culture flasks at 37 °C under a humidified atmosphere containing 5% CO_2_. Prior to experimental treatments, cells were serum-starved for 2 h in Dulbecco’s phosphate-buffered saline (DPBS) containing Ca^2+^ and Mg^2+^. Following serum starvation, cells were exposed to defined experimental conditions, including treatment with 3 µM cannabidiol (CBD), ionizing radiation at a exposure dose of 10 Gy, or a combination of CBD treatment followed by irradiation, as schematically illustrated in Figure 1. After each step lipid droplets were isolated. To validate lipid droplet isolation and enable selective tracking of lipid droplet–associated lipids, cells were additionally labeled with 10 µM d_31_-palmitic acid. The concentration of the deuterated fatty acid was selected based on MTS viability assays performed across a range of d^31^-palmitic acid concentrations, which identified 10 µM as a non-cytotoxic condition, while higher concentrations induced reduced cell viability (Figure 2). Incorporation of the deuterated fatty acid allowed lipid droplets to be identified via the characteristic C–D vibrational band in Raman spectra (Figure 2).

### CBD incubation, X-ray irradiation and combination

A CBD concentration of 3 µM, ionizing radiation at a exposure dose of 10 Gy, and their combined application were selected based on conditions previously established and characterized in our earlier studies investigating lipid metabolism, where cellular viability and DNA damage were systematically assessed using MTS and comet assays. [ACS sensors]

### Lipid droplets isolation

Lipid droplets were isolated from cultured cells using a gradient-based centrifugation approach according to the manufacturer’s protocol (Lipid Droplet Isolation Kit, ab242290, Abcam), with minor adaptations. Briefly, cells were harvested, washed with phosphate-buffered saline, and resuspended in Reagent A, followed by incubation on ice to facilitate cell lysis. Subsequently, Reagent B was added and samples were further incubated on ice prior to mechanical homogenization. Cell homogenates were gently layered with Reagent B to form a density gradient and centrifuged at 20,000 × g for 3 h at 4 °C. Following centrifugation, the lipid droplet–enriched fraction was recovered from the top of the gradient by careful pipetting and transferred to fresh microcentrifuge tubes. Isolated lipid droplets were either used immediately for Raman spectroscopic analysis or stored at −80 °C until further use, as recommended. To exclude possible contamination of iLDs fraction with buffer residues, Raman spectra for both reagents were recorded (Figure 5)

### Spectroscopic investigations

Raman spectroscopic measurements were performed on isolated lipid droplets (iLDs) deposited onto calcium fluoride (CaF_2_) windows (Crystran Ltd., UK). Briefly, 10 µL of lipid droplet suspension in isolation buffer was deposited onto the substrate and allowed to settle for approximately 15 min. Subsequently, samples were gently rinsed three times with ultrapure water to remove residual buffer components and immediately dried under a gentle nitrogen stream.

Raman spectra of iLDs were acquired using a Renishaw InVia Raman spectrometer coupled to a confocal optical microscope. Excitation was provided by an air-cooled solid-state laser operating at 532 nm, and the scattered signal was detected using a CCD detector maintained at −70 °C. A 100× air objective was used to localize individual lipid droplets and collect single-point Raman spectra. Spectra were recorded at 50% laser power with an integration time of 10 s and a single accumulation.

Raman imaging was performed on the cytoplasmic regions of cells in PBS buffer, labeled with d_31_-palmitic acid. Raman maps were acquired for 10 individual cells using a step size of 0.5 µm, an integration time of 0.3 s per pixel (one accumulation), and a spectral resolution of approximately 1.5 cm^−1^ with 60x immersive objective.

Raman images and single-point spectra were processed using WiRE software (version 5.3, Renishaw, United Kingdom). Data preprocessing included cosmic ray removal, noise filtering, baseline correction, and min–max normalization of all spectra. Hierarchical cluster analysis (HCA) was subsequently applied to identify spectral classes corresponding to lipid droplets accumulating d_31_-palmitic acid. Spectral distances were calculated using Euclidean distance, and clusters were defined using Ward’s linkage method.

Principal component analysis (PCA) was performed using Unscrambler X software (version 10.3). PCA was calculated for selected spectral regions using the non-linear iterative partial least squares (NIPALS) algorithm with leave-one-out cross-validation. The initial decomposition included seven principal components with 20 iterations, and the results were visualized as three-dimensional score plots accompanied by corresponding loading plots.

Integrated intensities were calculated for selected Raman bands associated with lipid chain conformation and packing (1060, 1131, 1167, and 1292 cm^−1^). All analyses were performed at the level of individual spectra to account for spectral heterogeneity and unequal numbers of spectra across experimental conditions. For each Raman band and experimental condition, median integrated intensities were calculated and used to determine fold changes relative to untreated controls. Fold changes were log_2_-transformed and visualized as heatmaps, with rows representing Raman bands and columns representing experimental conditions. The color scale was centered at zero, such that positive and negative values indicate relative increases or decreases relative to control samples. Identical color scale limits were applied across all heatmaps to enable direct comparison between cell types and treatments.

## Authors contribution

**Karolina Chrabąszcz:** conceptualization, data curation, formal analysis, funding acquisition, investigation, methodology, project administration, resources, supervision, validation, visualization, writing–original draft, writing–review and editing.

**Agnieszka Panek:** investigation, methodology, writing–review and editing

**Katarzyna Pogoda:** conceptualization, methodology, supervision, writing–review and editing

## Acknowledgments

This work was supported by the National Science Centre, Poland (2023/51/D/ST4/01686).

